# HIF1A-AS2 Regulates Hypoxia-Driven HIF Isoform Switching in Human Endothelial Cells

**DOI:** 10.64898/2026.11.06.764826

**Authors:** Anna Barton, Anna Więch -Walów, Paulina Czechowicz, Aleksandra Cabaj, Liliana Schaefer, James F. Collawn, Rafal Bartoszewski

**Author notes:** Correspondence: Rafal Bartoszewski, Department of Biophysics, Faculty of Biotechnology, University of Wroclaw, F. Joliot-Curie 14a Street, Wroclaw, 50-383, Poland.

## Abstract

Hypoxia-inducible factors (HIFs) are central regulators of cellular adaptation to low oxygen, with a well-documented transition from HIF-1α to HIF-2α signaling during prolonged hypoxia. Here, we investigate the role of a natural antisense transcript of *HIF1A*, *HIF1A-AS2*, in modulating this transition in human endothelial cells (ECs). Genome-wide expression profiling revealed a strong hypoxia-induced upregulation of *HIF1A-AS2* that coincided with a marked reduction in *HIF1A* mRNA levels, whereas *EPAS1* remained stable. Interestingly, this negative correlation was consistent across multiple EC types and supported by silencing experiments which demonstrated that both HIF-1α and HIF-2α contribute to *HIF1A-AS2* induction. Knockdown of *HIF1A-AS2* partially restored *HIF1A* expression and increased HIF-1α protein and functional activity, as evidenced by elevated *VEGFA* and *GLUT1* expression. Genomic analysis identified HIF-1 and HIF-2 binding motifs upstream of *HIF1A-AS2*, supporting the dual regulation. Furthermore, cancer tissue analysis revealed frequent downregulation of *HIF1A-AS2*, correlating with elevated *HIF1A* and *EPAS1* levels. These findings suggest that *HIF1A-AS2* acts as a hypoxia-responsive regulator that limits HIF-1α signaling and facilitates the HIF isoform switch, and suggests implications for endothelial adaptation, angiogenic signaling, and vascular remodeling under chronic hypoxia.

## Introduction

Hypoxia-inducible factors (HIFs) enable cells to survive and restore oxygen homeostasis when the demand for oxygen is not met—a condition known as hypoxia [1–3]. The activity of these transcription factors allows cells to extensively reprogram their gene expression profiles to favor glycolysis, stimulate angiogenesis, and promote erythropoietin production [1,4–6].

Transcriptionally active HIFs are heterodimers composed of α- and β-subunits that assemble during hypoxia, when the α-subunit is dramatically stabilized [7,3]. When cellular oxygen demands are met, HIF-α subunits undergo prolyl hydroxylation by specific prolyl hydroxylases (PHDs), which targets them for rapid proteasomal degradation [8,9]. Since PHD activity is oxygen-dependent, these enzymes become inactive under hypoxic conditions, allowing HIF-α subunits to accumulate and heterodimerize with β-subunits. The resulting transcriptionally active HIF complexes translocate to the nucleus, where they interact with hypoxia-response elements (HREs) in the promoters and enhancers of numerous target genes [3,10,11].

Importantly, prolonged or excessive adaptation to hypoxic conditions may endanger cells upon the rapid restoration of oxygen levels, potentially leading to oxidative damage [12,4]. Therefore, the extent of HIF-mediated adaptation must be carefully controlled to ensure safe recovery from hypoxic stress [13,14].

In specific cell types, including endothelial cells (ECs), cardiomyocytes, and hepatocytes as well as in many cancer cells, prolonged hypoxia is accompanied by a gradual transition between two HIF-α isoforms: HIF-1α, which is present in all cell types, and HIF-2α, which is tissue-specific [14–16]. This so-called “HIF switch” enables both normal and cancer cells to adjust transcriptional reprogramming in response to prolonged oxygen deficiency, ensuring survival while minimizing the risk of oxidative damage [14]. Thus, understanding the mechanisms underlying the HIF isoform transition may be crucial for the development of novel therapies for various human diseases. In endothelial cells, the HIF isoform switch is a critical determinant of angiogenic competence, metabolic adaptation, and survival during prolonged hypoxia.

Importantly, the HIF transition in ECs is a consequence of a decline in mRNA encoding HIF-1α (*HIF1A*) and the more stable levels of the HIF-2α transcript (*EPAS1*) [14,7]. This, combined with glycolysis-related reinstatement of PHD activity, leads to faster elimination of HIF-1α [7]. Although several mechanisms have been proposed to explain the destabilization of *HIF1A* mRNA during hypoxia, including the involvement of microRNAs and long non-coding RNAs, their significance in the HIF switch remains poorly understood [14,17–20].

Among these, long non-coding RNAs that act as natural antisense transcripts (NATs) of *HIF1A* (*HIF1A-AS1*, *HIF1A-AS2*, *HIF1A-AS3* aka *HIF1AL*, and *HIF1AS*) have been shown to reduce HIF-1α transcript levels, often functioning in a negative feedback loop with HIF-1 [19,21–23].

In this study, we observed that *HIF1A-AS2* levels are elevated under prolonged hypoxia in various types of human ECs and negatively correlate with *HIF1A* expression. Furthermore, we demonstrate that this antisense transcript is regulated not only by HIF-1 but also by HIF-2, ensuring its expression during prolonged hypoxia even in the absence of HIF-1α. Notably, although *HIF1A-AS2* significantly impacts hypoxic *HIF1A* mRNA levels, its silencing was not sufficient to restore HIF-1α levels under prolonged hypoxia, suggesting that it plays a complemenary role with microRNAs in the hypoxic HIF transition.

## Results

To initially access potential influence of natural antisense transcripts of *HIF1A* on the transition from HIF-1α to HIF-2α in human ECs (**Figure 1A**), we analyzed their expression alongside with *HIF1A* and *EPAS1* mRNA levels using genome-wide microarray data obtained from Human Umbilical Vein ECs (HUVECs; 3 biological replicates, each comprising a pool of 10-donor each [15]) following 2 hour, 8 hour and 16 hour exposure to hypoxia (0.9% O_2_) (**Figure 1**).

**Figure 1.**
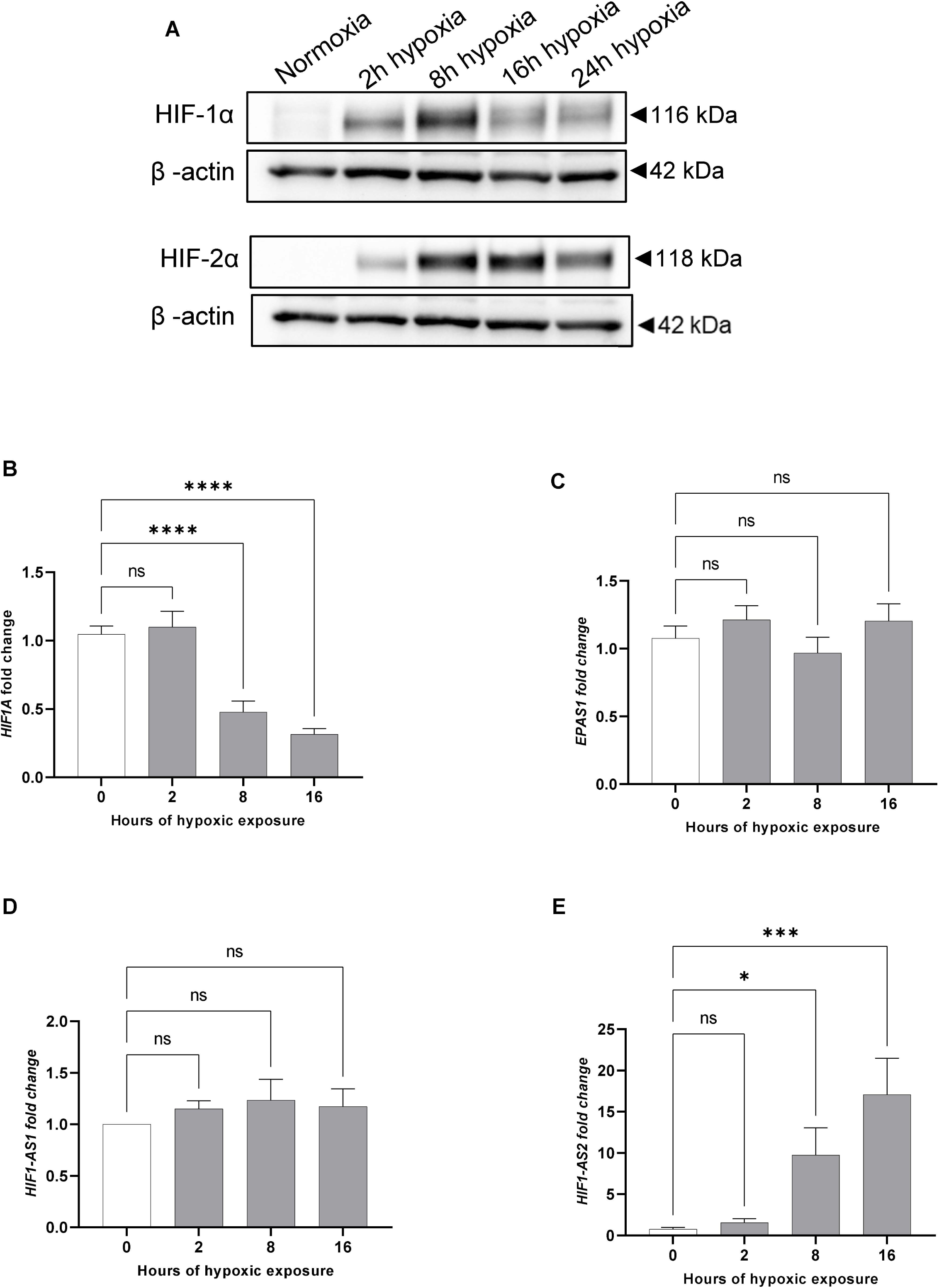
The HIF switch is accompanied by negatively correlated changes in HIF1A and HIF1A-AS2 mRNA levels in HUVECs. (**A**) Representative changes in HIF-1α and HIF-2α protein levels were evaluated by Western Blot after exposure to hypoxia for the time periods specified. Uncropped blots are presented in **Supplemental** Figure 2. The corresponding changes in *HIF1A* (**B**), *EPAS1*(***C***), *HIF1-A*S1(**D**) *HIF1-A*S2(**E**) mRNA levels were quantified based on expression microarray signals and expressed as fold change relative to normoxic conditions. The data represents mean of 3 biological replicates ± SEM of (\**P*<0.05, \*\**P*<0.01 and \*\*\**P*<0.001 were considered significant).

Consistent with our previous findings, *HIF1A* transcript levels were dramatically reduced starting at the 8-hour time point (**Figure 1B**), whereas *EPAS1* mRNA levels remained stable until 16 hours (**Figure 1C**) [15]. This data corresponded well with the accumulation patterns of related HIF-α subunits during hypoxia: HIF-1α levels were significantly reduced after 16 h while HIF-2α protein remained elevated until 24 hours (**Figure 1A**). Notably, while the expression of *HIF1A-AS1* (NR_047116.1) remained unaffected by hypoxia (**Figure 1D**), the *HIF1A-AS2* (NR_045406.1) was hypoxia-inducible, showing an approximately 8-fold increase at 8 hours, and doubling again by 16 hours (**Figure 1E**). The probes for the *HIF1A-AS3/HIFAL* (<u>NR_144368.1</u>) [23] and the *HIF1A-AS* [19] transcript were not present on the microarray platform at the time of analysis. Nevertheless, the observed negative correlation between *HIF1A* mRNA and *HIF1A-AS2* expression suggested that this NAT may contribute to the destabilization of *HIF1A* transcript in ECs.

To determine whether *HIF1A-AS2* expression is associated with the reduction of *HIF1A* transcript levels in other endothelial cell types, we conducted parallel hypoxia studies in eight human primary endothelial cell lines derived from different vascular beds, as well as immortalized HUVECs (HUVEC hTERT) (**Figure 2**). This analysis revealed that *HIF1A-AS2* levels in HUVECs increased rapidly after 8 hours of hypoxia exposure, and a similar pattern was observed in most of the tested cell lines. However, the magnitude of *HIF1A-AS2* induction was cell type-specific, ranging from a 6-fold increase at 16 hours in Human Aortic Endothelial Cells (HAEC) and Human Pulmonary Artery ECs (HPAEC) (**Figure 2DF**), respectively, to a 40- to 60-fold increase in Human Lung Microvascular ECs (HMVEC-L) and Human Coronary Artery ECs HCAEC (**Figure 2EJ**).

**Figure 2.**
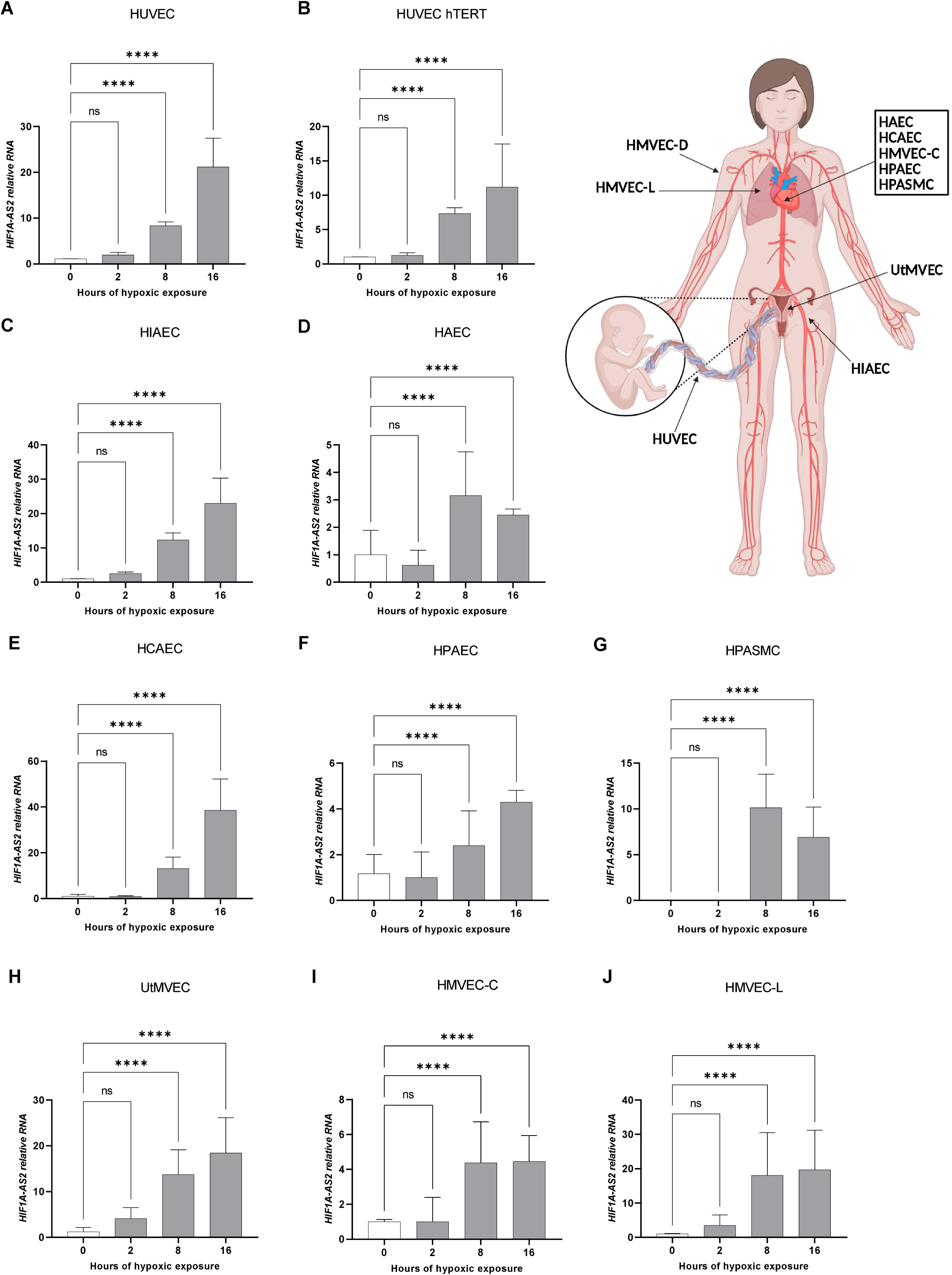
Prolonged hypoxia results in the accumulation of HIF1A-AS2 RNA in human endothelial cells. HUVEC (**A**), HUVEC hTERT (**B**), HIAEC (**C**), HAEC (**D**), HCAEC (**E**); HPAEC (**F**), HPASMC (**G**), UtMVEC (**H**), HMVEC-C (**I**) and HMVEC-L (**J**), cells were exposed to hypoxia for the time periods specified and total RNA was collected The corresponding *HIF1A*-*AS2* RNA levels were quantified by quantitative real-time PCR and normalized to *RPLP0* rRNA levels and expressed as a fold change over control at the specific time-point of hypoxia. Data represent the mean ± SEM of three replicates (\**P*<0.05, \*\**P*<0.01 and \*\*\**P*<0.001 were considered significant). The diagram depicting ECs locations in Human was created in https://BioRender.com.

Consistent with previous reports [15,24], *HIF1A* mRNA levels measured in parallel were significantly reduced in all tested endothelial cell types starting from 8 hours of hypoxia (**Figure 3**). Taken together, these results demonstrate a strong negative correlation between *HIF1A* and *HIF1A-AS2* expression during the hypoxic time course across all tested endothelial cell lines, as confirmed by statistical analysis of their expression profiles (**Figure 3K**).

**Figure 3.**
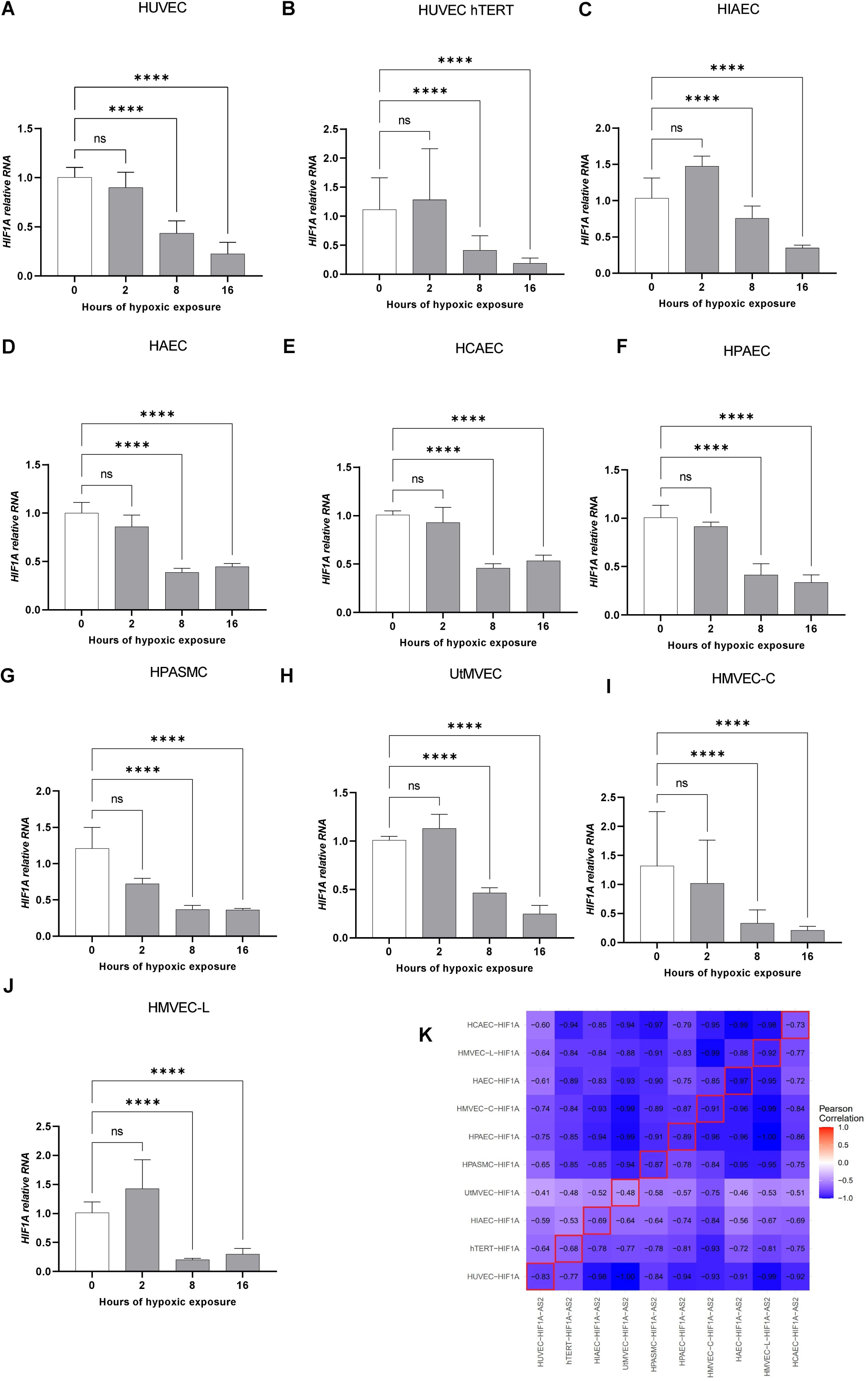
Prolonged hypoxia results in the accumulation of HIF1A mRNA in human endothelial cells. HUVEC (**A**), HUVEC hTERT (**B**), HIAEC (**C**), HAEC (**D**), HCAEC (**E**); HPAEC (**F**), HPASMC (**G**), UtMVEC (**H**), HMVEC-C (**I**) and HMVEC-L (**J**), cells were exposed to hypoxia for the time periods specified and total RNA was collected The corresponding *HIF1A*-*AS2* RNA levels were quantified by quantitative real-time PCR and normalized to *RPLP0* rRNA levels and expressed as a fold change over control at the specific time-point of hypoxia. Data represent the mean ± SEM of three replicates (\**P*<0.05, \*\**P*<0.01 and \*\*\**P*<0.001 were considered significant). (**K**) Pearson correlation analysis of time dependent expression of *HIF1A-AS2* and *HIF1A* mRNA in ECs.

Next, to gain insight into the mechanism responsible for hypoxic induction of *HIF1A-AS2*, we examined the genomic location of this NAT within open chromatin regions identified in the HUVEC cell line by the ENCODE project for the presence of distinct HRE motifs for HIF-1 and for HIF-2 [25–27] (**Figure 4**). Consistent with previous reports suggesting a feedback loop between HIF-1 and *HIF1A-AS2* [*28,29*], we identified two HIF-1 binding motifs upstream of the *HIF1A-AS2* transcription start site (TSS). Notably, we also found a HIF-2 binding motif in this region (**Figure 4C**). All these motifs are located relatively close to TSS and to each other, suggesting a potentially coordinated regulatory mechanism (**Figure 4C**). In contrast, although *HIF1A-AS1* contains two HIF-1 motifs, only one is located upstream its TSS, and this NAT location lacks a HIF-2 motif entirely (**Figure 4B**). These findings suggest that the hypoxic induction of *HIF1A-AS2* may be mediated not only by HIF-1α but also by HIF-2α. This would be particularly important during prolonged hypoxia when HIF-2α becomes the dominant isoform. We hypothesize that HIF-2α contributes to the sustained expression of *HIF1A-AS2* and thereby prevents the re-accumulation of *HIF1A* mRNA when HIF-1α levels are low.

**Figure 4.**
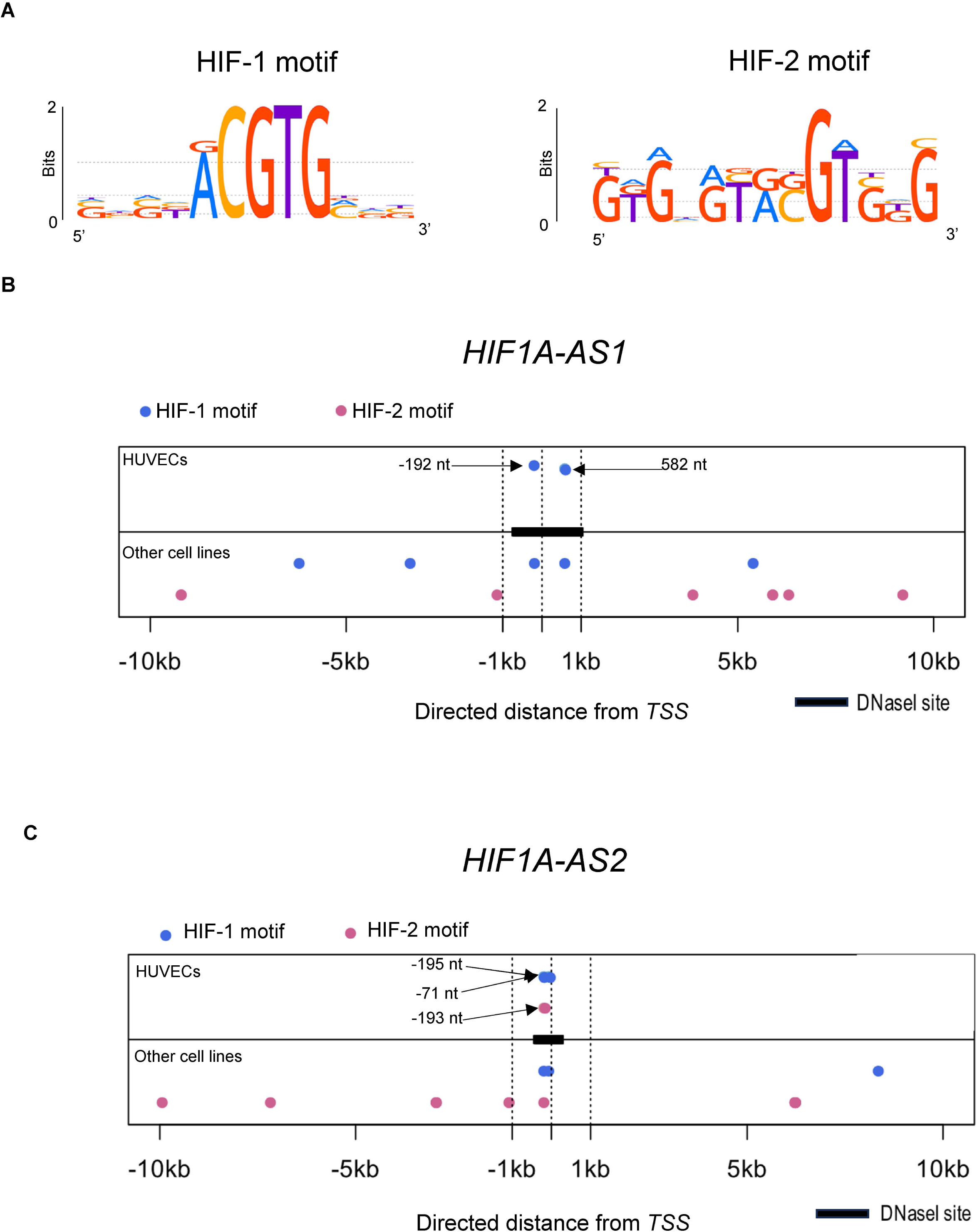
Differential HIF binding motif architecture suggests distinct regulation of HIF1A-AS1 and HIF1A-AS2. (**A**) Sequence logos of HIF-1 and HIF-2 hypoxia response element (HRE) binding motifs from HOCOMOCO v9 used in this analysis. (**B**) Graphical representation of HIF-1 (blue) HRE binding motifs in the *HIF1A-AS1* locus. (**C**) Graphical representation of HIF-1 (blue) and HIF-2 (orange) HRE binding motifs in the *HIF1A-AS2* locus, based on Nencki Genomics Database v79_1. Top panels show motif distribution in regions of open chromatin specific to HUVEC cells, while bottom panels represent motif distribution across all human cell lines included in the database.

To further test this hypothesis, we performed silencing experiments in HUVECs exposed to hypoxia for 8 hours (when both HIF-1α and HIF-2α are present) and 16 hours (when HIF-2α is dominant) (**Figure 5AB**). As shown in **Figure 5C**, silencing of *HIF1A* or *EPAS1* efficiently reduced *HIF1A-AS2* levels by approximately 50% at both time points, without affecting *HIF1A-AS1* expression. Moreover, simultaneous silencing of both *HIF1A* and *EPAS1* resulted in an even greater reduction of *HIF1A-AS2* levels, while again having no significant impact on *HIF1A-AS1* (**Figure 5D**). These results indicate that both HIF-1α and HIF-2α contribute to the induction of *HIF1A-AS2* during hypoxia.

**Figure 5.**
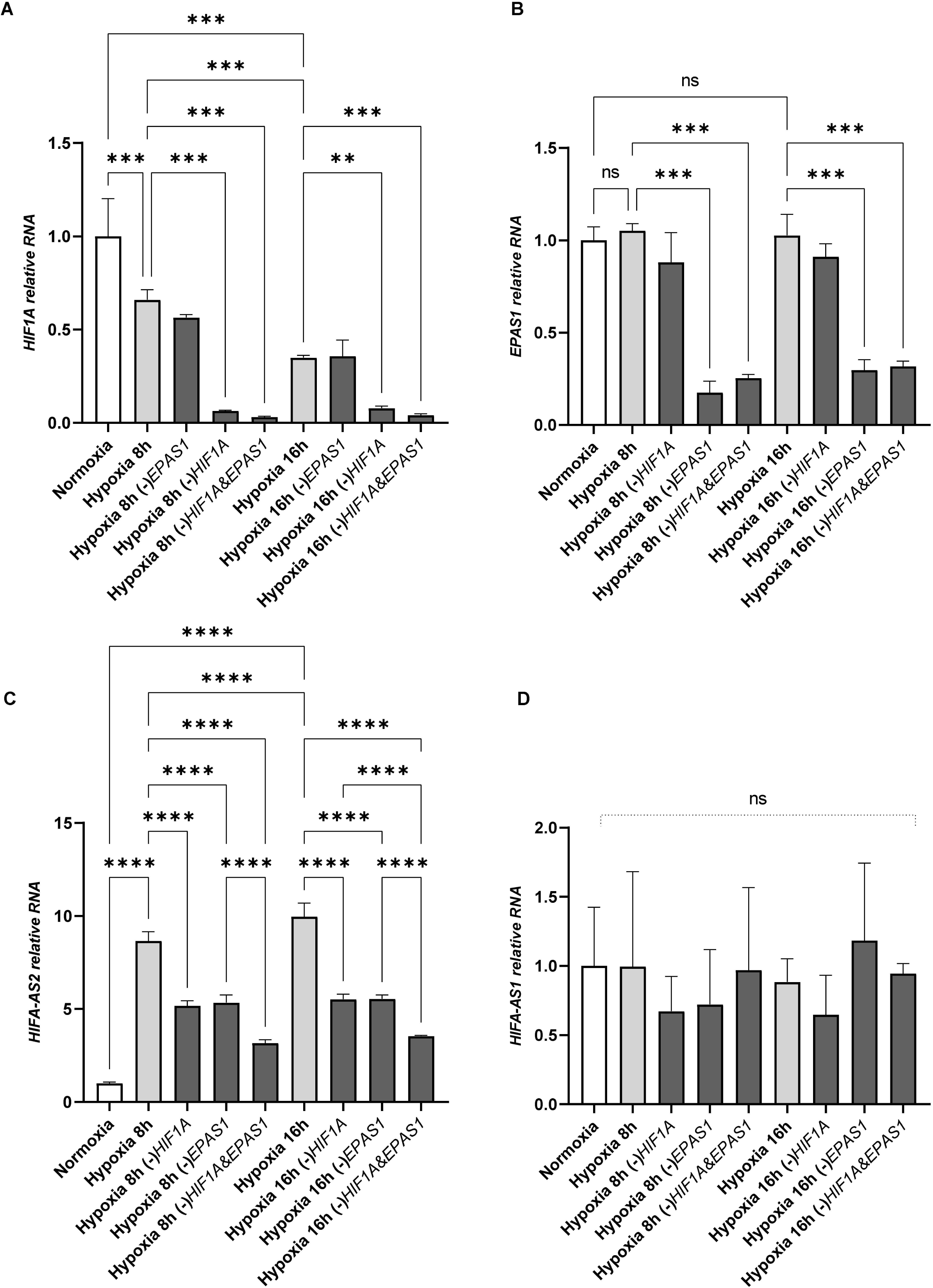
HIF1A and EPAS1 silencing in HUVECs exposed to hypoxia reduces the expression of HIF1A-AS2 but not HIF1A-AS1. The corresponding changes in HIF1A (**A**), EPAS1 (**B**), HIF1A-AS2 (**C**), HIF1A-AS1 (**D**) RNA levels were quantified by quantitative real-time PCR and normalized to RPLP0 rRNA levels and expressed as a fold change over control at the specific time-point of hypoxia. Data represent the mean ± SEM of three replicates (\**P*<0.05, \*\**P*<0.01 and \*\*\**P*<0.001 were considered significant).

Next, we examined the consequences of silencing *HIF1A-AS2* on hypoxic HIF-1α expression. As shown in **Figure 6A**, silencing *HIF1A-AS2* with specific siRNA reduced its RNA levels by approximately 50% in HUVECs exposed to hypoxia for 8 and 16 hours. Parallel assessment of *HIF1A* expression revealed that knockdown of *HIF1A-AS2* significantly increased *HIF1A* mRNA levels (**Figure 6B**). However, while *HIF1A-AS2* silencing led to increased *HIF1A* expression after 8 hours of hypoxia, the effect of siRNA on *HIF1A* transcript levels was significant but much less pronounced after 16 hours of exposure. Subsequent analysis of HIF-1α protein levels showed that reduction of *HIF1A-AS2* resulted in only a modest accumulation of HIF-1α after 8 hours of hypoxia, with no noticeable effect observed after 16 hours (**Figure 6C**). As these results were not sufficient to conclude that *HIF1A-AS2* has a functional effect on hypoxic HIF-1α accumulation, we next investigated whether the effects of this NAT on *HIF1A* expression would be reflected in HIF-1 activity. As shown in **Figures 6D** and **6E**, the hypoxic mRNA levels of vascular endothelial growth factor A (*VEGFA)* and glucose transporter 1 (*GLUT1*), canonical transcriptional targets of HIF-1, were significantly increased (by approximately 50%) upon *HIF1A-AS2* silencing after 8 hours of hypoxia, while after 16 hours, expression of *VEGFA* was unaffected, while *GLUT1* mRNA levels were only moderately increased. Taken together, these data suggest that, despite the modest effects of *HIF1A-AS2* on HIF-1α accumulation in hypoxia-exposed cells, this NAT efficiently limits HIF-1 activity and contributes to the transition from HIF-1 to HIF-2 signaling. However, accumulation of *HIF1A-AS2* during prolonged hypoxia may have limited effects on HIF-1α accumulation and related signaling.

**Figure 6.**
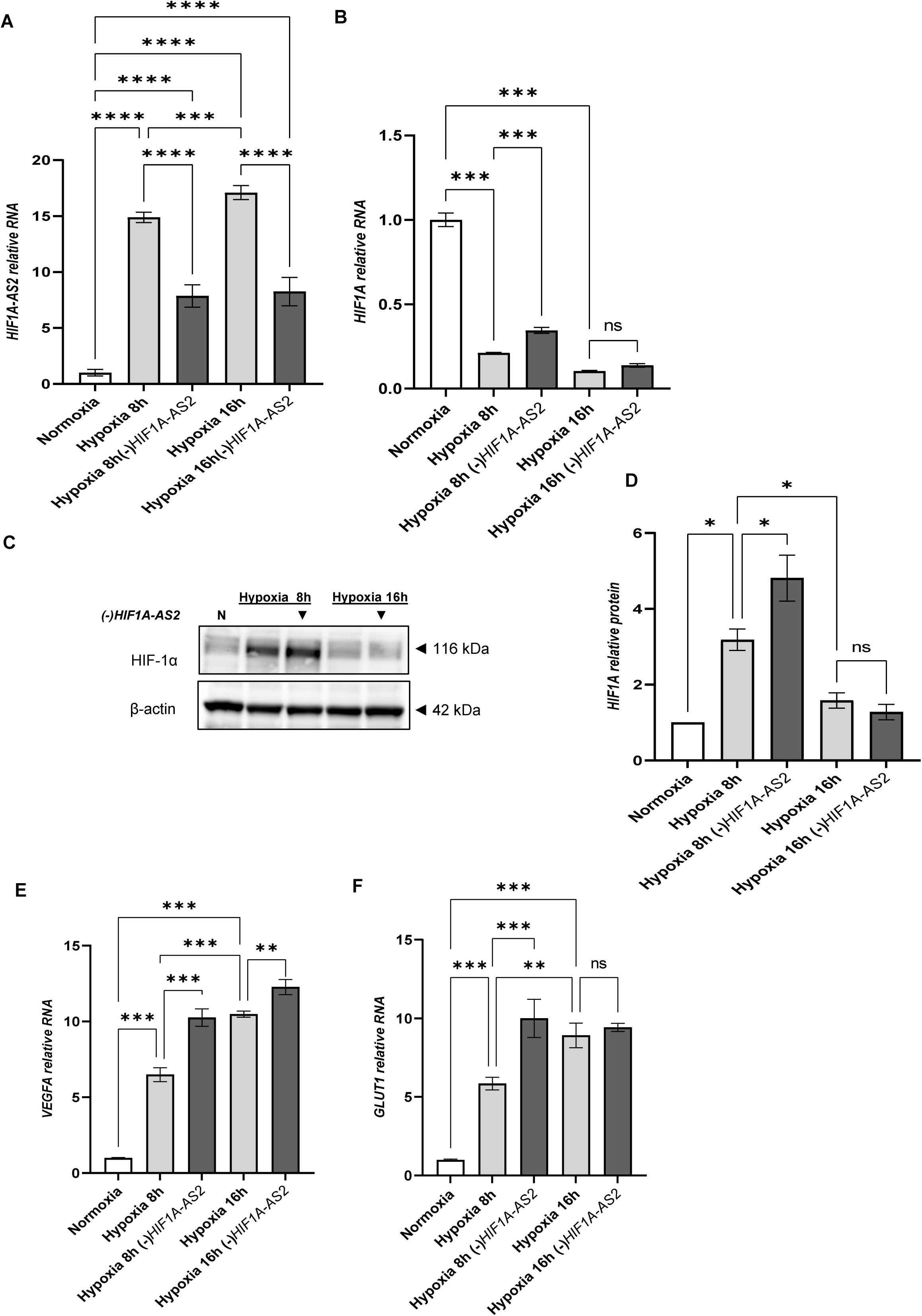
HIF1A-AS2 silencing in HUVECs exposed to hypoxia increases HIF-1α levels and downstream signaling. The corresponding changes in *HIF1A-AS2* (**A**) and *HIF1A* (**B**) RNA levels were quantified by quantitative real-time PCR and normalized to *RPLP0* rRNA levels and expressed as a fold change over control at the specific time-point of hypoxia. Data represent the mean ± SEM of three replicates (\**P*<0.05, \*\**P*<0.01 and \*\*\**P*<0.001 were considered significant). *HIF1A-AS2* silencing (depicted with ▾) related changes in HIF-1α protein levels were evaluated in normoxia (N) and after 8 h and 16 h exposure to hypoxia by Western Blot (**C**) and normalized to β-actin and total protein levels (**D**). Uncropped blots are presented in **Supplemental** Figure 2. The corresponding changes in *VEGFA* (**C**) and *GLUT1* (**D**) mRNA levels were quantified by quantitative real-time PCR and normalized to *RPLP0* rRNA levels and expressed as a fold change over control at the specific time-point of hypoxia. Data represent the mean ± SEM of three replicates (\**P*<0.05, \*\**P*<0.01 and \*\*\**P*<0.001 were considered significant).

Finally, to gain more translational insight into the relationship between *HIF1A* and its associated natural antisense transcripts, we analyzed data from the Gene Expression database of Normal and Tumor tissues 2 (GENT2) [30]. Specifically, we examined the expression levels of *HIF1A*, *EPAS1*, *HIF1A-AS1*, and *HIF1A-AS2* in both human normal and cancer tissues (**Figure 7A–E**).

**Figure 7.**
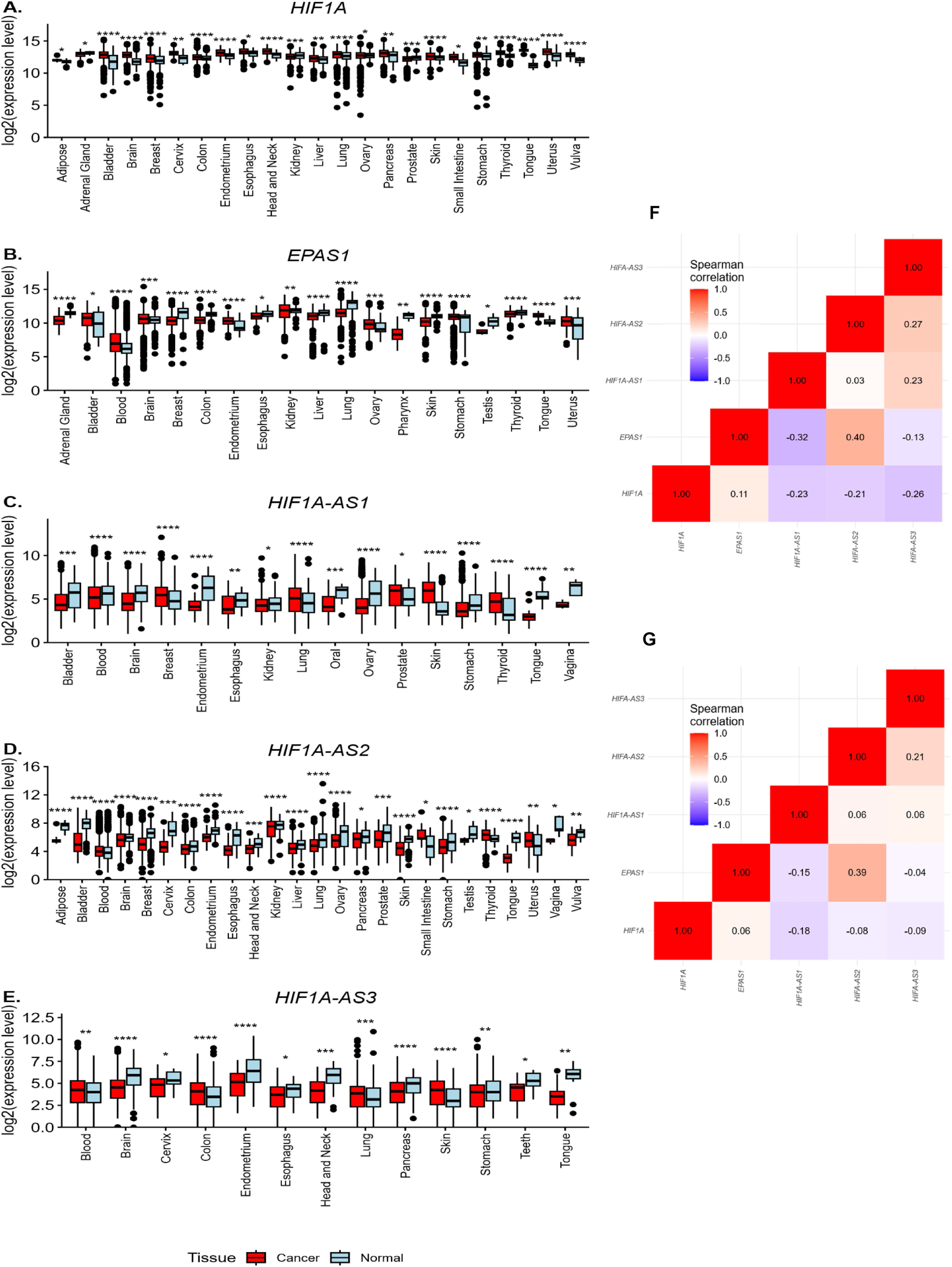
The expression of HIF1A-AS2 is negatively correlated with HIF1A mRNA and positively correlated with EPAS1 in normal tissues. These expression patterns, however, are deregulated in cancer cells. Data from the GENT2 were analyzed to assess the expression levels of HIF1A (**A**), EPAS1 (**B**), HIF1A-AS1 (**C**), HIF1A-AS2 (**D**), and HIF1A-AS3 (E) in both human normal (blue) and cancer tissues (red). The graphs include only those tissue pairs that showed statistically significant differences (*P <0.05, **P <0.01, ***P <0.001,****P <0.001). Spearman correlation analysis of the expression levels of HIF1A, EPAS1, HIF1A-AS1, HIF1A-AS2, and HIF1A-AS3 was performed separately for normal tissues (**F**) and tumors (**G**). This analysis was based on the complete GENT2 dataset, which includes expression data from all tissues, regardless of whether they showed statistically significant differences between normal and cancerous states.

As expected, *HIF1A* and *EPAS1* expression levels were significantly higher in the majority of cancer tissues compared to their normal counterparts (**Figure 7AB**), which is consistent with previously published data. Although all tested NATs showed some tendency toward deregulation in cancer cells, the differences in *HIF1A-AS1* and *HIF1A-AS3* expression between normal and cancer tissues were not particularly pronounced, despite occasional reductions in cancer samples (**Figure 7CE**).

Notably, however, as shown in **Figure 7D**, *HIF1A-AS2* expression was significantly reduced in many cancer tissues, suggesting a potential correlation with *HIF1A* expression. Follow-up statistical analyses (**Figures 7F–G**) confirmed this observation, revealing a negative correlation between *HIF1A-AS2* and *HIF1A* levels, particularly in normal tissues. However, this negative correlation was diminished in cancer cells, indicating that the reduced expression of this NAT may facilitate HIF-1-mediated hypoxic adaptation in these tissues. Furthermore, *HIF1A-AS2* expression showed the strongest positive correlation with *EPAS1* among all tested genes in both normal and cancer tissues, supporting our earlier findings that HIF-2 may primarily drive the accumulation of this NAT. Taken together, these results suggest that *HIF1A-AS2* may serve as an important regulator of the hypoxic HIF-switch in both normal and cancer tissues.

## Discussion

The comparatively lower stability of *HIF1A* mRNA relative to *EPAS1* has been identified as a key determinant in the transition from HIF-1α to HIF-2α in endothelial cells exposed to hypoxic conditions [7,15]. Under prolonged hypoxia, *HIF1A* transcript levels decline markedly, whereas *EPAS1* remains relatively stable [7,15]. This differential stability is likely due to *EPAS1’s* reduced susceptibility to AU-rich element (ARE)-mediated decay and resistance to miRNA-dependent degradation [15,31].

Importantly, chronic hypoxia impairs miRNA biogenesis via a von Hippel-Lindau (VHL)-dependent mechanism [32], which may attenuate miRNA-mediated repression of *HIF1A*. In this context, natural antisense transcripts such as *HIF1A-AS2* may contribute to the post-transcriptional downregulation of *HIF1A* [28,29], thereby reinforcing the HIF isoform switch and sustaining low *HIF1A* expression during prolonged hypoxic exposure. In the present study, we investigated the contribution of these regulatory mechanisms to the HIF-α isoform transition in endothelial cells.

Given the abundance of protein-coding loci overlapping with natural antisense transcripts in the human genome [33], yet the limited number shown to exert functional inverse regulation [34,19], we analyzed our previously published microarray dataset from hypoxia-exposed HUVECs derived from 30 donors, all of which exhibited a clear HIF isoform switch [15]. Due to the absence of specific probes for *HIF1A-AS3* [22] and the recently identified *HIF1-AS* [19] in the microarray platform used, we were able to assess expression changes for only two *HIF1A*-specific NATs: *HIF1A-AS1* and *HIF1A-AS2*.

Independent analysis of next-generation sequencing data from the same cell population indicated that *HIF1A-AS3* is not expressed in these cells [24]. However, the lack of data on *HIF1-AS* represents a limitation of our study. This transcript was only recently identified in endothelial cells through *de novo* transcriptome assembly [19], and to date, no specific TaqMan probes are available to validate its expression during hypoxic time courses.

Importantly, *HIF1-AS* has been shown to remain in the nucleus, where it modulates histone H3 methylation at the *HIF1A* promoter, thereby repressing *HIF1A* pre-mRNA expression [19]. This nuclear mechanism contrasts with mRNA destabilization processes that typically occur in the cytoplasm. Since this pathway primarily influences transcriptional initiation rather than mRNA stability, *HIF1-AS* likely functions through a distinct regulatory mechanism and is therefore considered beyond the scope of the present study.

Using an independent TaqMan probe-based qPCR approach across multiple primary endothelial cell types, we confirmed that the convergent antisense transcript *HIF1A-AS2*, which partially overlaps the 3′ untranslated region (UTR) of *HIF1A* [35], is significantly and consistently upregulated during hypoxia, reaching peak expression under chronic conditions. Although earlier *in silico* analyses suggested that *HIF1A-AS2* may interact with AU-rich elements within the *HIF1A* 3′UTR to promote transcript destabilization [28], this mechanism remains under debate [36,23].

Paradoxically, *HIF1A-AS2* has previously been also proposed to function as a molecular sponge for miR-153-3p in HUVECs, thereby enhancing *HIF1A* expression and promoting HIF-1α accumulation under hypoxic conditions [37]. However, our data do not support this model. Furthermore, during our previous extensive research on hypoxia-related changes in miRNA levels in HUVECs we did not observed any significant changes in hsa-miR-153-3p expression [24]. Instead, our findings clearly demonstrate that *HIF1A-AS2* contributes to the downregulation of *HIF1A* expression in HUVECs, supporting its role in facilitating the HIF isoform switch during prolonged hypoxia. This conclusion is further substantiated by a consistent negative correlation between *HIF1A-AS2* and *HIF1A* mRNA levels observed across endothelial cells derived from various vascular beds under hypoxic conditions. This suggests that *HIF1A-AS2* may contribute to the destabilization of *HIF1A* transcripts, potentially facilitating the transition from HIF-1α to HIF-2α signaling.

Our study shows that the hypoxic expression of *HIF1A-AS2* is part of a negative-feedback system. Mechanistically, the presence of both HIF-1 and HIF-2 binding motifs upstream of *HIF1A-AS2* supports a dual regulatory model, where initial induction is mediated by HIF-1α, and sustained expression during prolonged hypoxia is maintained by HIF-2α. Silencing experiments further confirmed that both transcription factors contribute to *HIF1A-AS2* expression, reinforcing the hypothesis of a feedback loop that limits HIF-1α activity.

Functionally, knockdown of *HIF1A-AS2* resulted in increased *HIF1A* mRNA levels and a modest elevation of HIF-1α protein, particularly at early hypoxic time points (e.g., 8 hours). Although the effect on HIF-1α protein accumulation was limited, the observed upregulation of canonical HIF-1 target genes, such as *VEGFA* and *GLUT1*, following *HIF1A-AS2* silencing suggests that this NAT contributes to the fine-tuning of HIF-1 transcriptional activity.

It is important to note that our ability to effectively silence *HIF1A-AS2* in HUVECs was limited. Despite optimization of transfection protocols and the use of two consecutive transfection rounds, we achieved only approximately a 50% reduction in hypoxia-induced *HIF1A-AS2* expression with levels that remained above those observed under normoxic conditions. Consequently, we cannot exclude the possibility that *HIF1A-AS2* may exert a more substantial effect on HIF-1α levels and downstream signaling under conditions of a more efficient knockdown. This highlights the potential of *HIF1A-AS2* as a regulatory target for restoring HIF-1α activity during prolonged hypoxia.

HIF-1α accumulation during chronic hypoxia is also restricted by several independent mechanisms, including miRNA-mediated repression and transcriptional inhibition [14,20,38,17]. Therefore, we propose that *HIF1A-AS2* primarily functions as a negative regulator of HIF-1 signaling during the early phases of hypoxia. Its regulatory influence appears to diminish over time, suggesting a role in fine-tuning the hypoxic response and potentially contributing to the transition from HIF-1α to HIF-2α dominance.

Previous studies have strongly associated *HIF1A-AS2* with various malignancies, including non-papillary renal cell carcinoma, breast cancer, and glioblastoma [35,39,40,23], as well as ischemic conditions such as myocardial infarction [41,42]. To further explore its relevance, we conducted a comprehensive analysis of *HIF1A-AS2* expression across normal and cancerous tissues. This analysis revealed that *HIF1A-AS2* is frequently downregulated in tumors, despite elevated expression of *HIF1A* and *EPAS1*. These findings suggest that loss of *HIF1A-AS2* may contribute to sustained HIF-1α signaling in cancer, potentially promoting angiogenesis and metabolic adaptation.

Interestingly, we also observed a strong positive correlation between *HIF1A-AS2* and *EPAS1* expression in both normal and cancer cells. This finding not only supports the involvement of HIF-2α in the regulation of *HIF1A-AS2*, but also underscores the potential role of this NAT in modulating the HIF isoform switch in pathological contexts. Furthermore, silencing of the third HIF-α isoform, HIF-3α, which is known to accumulate during prolonged hypoxia and acts to suppress HIF-1α activity while promoting endothelial cell survival [43–45] resulted in a significant increase in *HIF1A-AS2* expression (**Supplementary Figure 1**). This observation suggests that cells may have evolved complementary regulatory mechanisms to finely tune HIF-1α signaling.

Taken together, our study provides new insights into the regulatory role of *HIF1A-AS2* in modulating the hypoxic response and facilitating the transition from HIF-1α to HIF-2α signaling in human endothelial cells. The frequent downregulation of *HIF1A-AS2* in tumors despite the elevated *HIF1A* and *EPAS* expression suggests a role in sustaining HIF-1α signaling in cancer. Its positive correlation with *EPAS1* and upregulation following HIF-3α silencing further support its broader involvement in coordinating HIF signaling. These findings support a model in which HIF1A-AS2 contributes to the temporal coordination of HIF signaling in endothelial cells, rather than acting as a binary on/off regulator.

## Materials and Methods

### Cell culture

Primary human umbilical vein endothelial cells (HUVEC) were purchased from Cellworks (Caltag Medsystems Ltd, UK) and cultured in EGM-2 Bulletkit Medium (Lonza). Primary human aortic endothelial cells (HAEC), primary human cardiac artery endothelial cells (HCAEC), primary human iliac endothelial cells (HIAEC), and primary human pulmonary artery endothelial cells (HPAEC) were purchased from Lonza and cultured in EGM-2 Bulletkit medium. Primary human dermal microvascular endothelial cells (HMVEC-D), primary human lung microvascular endothelial cells (HMVEC-L), primary human uterine microvascular endothelial cells (UtMVEC) were purchased from Lonza and cultured in EGM-2MV medium (Lonza). Primary human pulmonary artery smooth muscle cells (HPASMC) were purchased from Gibco and cultured in Medium 231 supplemented with SMGS (Gibco). Except for HUVECs that were pooled from ten individual donors, all other primary human ECs were obtained from single donors. Immortalized human umbilical vein endothelial cells HUVEC/TERT 2 - CRL-4053™ (ATCC, USA) were cultured in EGM-2 Bulletkit Medium. All primary cell culture experiments were conducted between passages 2 and 6 at a confluence of 70-80 %.

### Induction of hypoxia

Hypoxia was induced in a CO_2_/O_2_ incubator/chamber specific for **hypoxia** research (InvivO2 Baker Ruskinn). Briefly, cells were cultured in 35 mm dishes for RNA isolation and protein isolation, at 0.9% O_2_ for the time periods specified (PO_2_ was 6.84 mm Hg). Control cells were maintained in normoxia in a CO_2_ incubator (Thermo Fisher Scientific).

### siRNA transfections

siRNA for *HIF1A* (Silencer Select, Invitrogen, assay id s6539), *EPAS1* (Silencer Select, Invitrogen, assay id s4698), *HIF1A-AS1* (Silencer Select, Invitrogen, assay id n513072) and *HIF1A-AS2* (Silencer Select, Invitrogen, assay id n349857) were purchased from Invitrogen. HUVECs were transfected using the Lipofectamine RNAiMax (Invitrogen, 13778030) according to manufacturer’s protocol. The siRNAs were used at a final concentration of 20 nM (for *HIF1A*, *EPAS1* and *HIF1A-AS1*) and 40 nM for *HIF1A-AS2*. Silencer Select Negative Control #1 siRNA (Invitrogen, 4390843) was used as a control at a final concentration of 10 nM. Following transfection, the cells were cultured for 48 h prior to analysis, with the medium replaced 24 h post-transfection. After 48 h, cells were placed in a hypoxia chamber, whereas control cells were maintained under normoxic conditions.

### Isolation of RNA

The cells were lysed directly on culture vessels with RLT lysis reagent (Qiagen). Total RNA (containing both mRNA and miRNA) was isolated using miRNeasy kit (Qiagen). RNA concentrations were calculated based on the absorbance at 260 nm. RNA samples were stored at -70°C until use.

### Measurement of mRNA using quantitative Real Time PCR (qRT-PCR)

TaqMan One-Step RT-PCR Master Mix Reagents (Applied Biosystems) was used as described previously [38,46–49] using the manufacturer’s protocol. The relative expressions were calculated using the 2^-ΔΔCt^ method [50] with the Ribosomal Protein Lateral Stalk Subunit P0 (*RPLP0*) gene as reference genes for the mRNA. TaqMan probe ids used are given in the **Supplemental Table 1.**

### Western Blots

Western Blot analysis was performed as previously described [49]. Briefly, cells were lysed in RIPA Lysis and Extraction Buffer (Thermo Fisher Scientific) supplemented with protease inhibitors (cOmplete ™ Mini, Roche). The cells were homogenized by passing the lysate 20 times through a 0.4 mm needle attached to a sterile plastic syringe. The insoluble material was removed by centrifugation at 15,000g for 15 min. Protein concentrations were determined by Bio-Rad™ DC-Protein Assay using bovine serum albumin (BSA) as standard. Following the normalization of the protein concentrations, the lysates were mixed with an equal volume of 5X Laemmli sample buffer (10% SDS, 50% glycerol, 0.05% bromophenol blue, 12.5 % β-mercaptoethanol. 312,5 mM Tris-HCl, pH = 6.8) and incubated for 10 min at 95°C prior to separation by SDS-PAGE on a 4-15% Criterion TGX Stain-Free Gel (Bio-Rad, Hercules, CA, USA). PageRuler™ Plus Prestained Protein Ladder (Thermo Scientific, #26619) was used for molecular weight estimation. Following SDS-PAGE, the proteins were transferred to polyvinylidene fluoride membranes (Bio-Rad) using the wet electroblotting method (300 mA, 4°C, 90 min for one gel and 180 min for two gels). The membranes were blocked with BSA dissolved in TBS/Tween-20 (5% BSA, 0.1% Tween-20 overnight at 4°C), followed by immunoblotting with the primary antibodies (2 hours, RT) that included rabbit-anti-HIF-1a D1S7W (1:1000, #36169; Cell Signaling), rabbit anti-HIF-2a D6T8V (1:1000, #59973: Cell Signaling), rabbit anti-actin (1:1000, #4967; Cell Signaling). After the washing steps, the membranes were incubated with horseradish peroxidase–conjugated Goat Anti-Rabbit IgG (H+L) secondary antibodies (1:5000; #1706515, Bio-Rad) for 1 h at room temperature and detected using SuperSignal West Pico ECL (Thermo Scientific). Densitometry was performed using the Image Lab software v.4.1 (Bio-Rad).

### Hypoxia Response Element (HRE) analysis

The promoters of the miRNAs were analyzed for HIF-1 or HIF-2 binding sites. In each promoter sequence that was defined as a 20kb window around the transcription start site (TSS), only the open chromatin regions were examined that were established in the HUVEC cell line by the ENCODE [25] project. Both DNase I-seq HUVEC datasets found in Ensemb l were merged [51] (v.79). The focus was on two distinct HRE motifs annotated to HIF-1 (M00139, alias *HIF1A)* and HIF-2 (M00074, alias *EPAS1)* from the HOCOMOCO v.9 database [26]. We note that from the version 10 onwards, the creators of HOCOMOCO have decided to choose a single generic model for each TF [26], resulting in assignment of nearly identical motifs to HIF1A and *EPAS1*, however, based on previous research reports [52–55], we continue to use the two distinct HRE motifs annotated to *HIF1A* and *EPAS1*. The Nencki Genomics Database [56] (v. 79_1) was used to obtain genomic coordinates of these motif instances.

### Statistical analysis

Results were expressed as a mean ± standard deviation (SD). Statistical significance was determined using ANOVA and a *P* value <0.05 was considered significant. The correlations were accessed via the Pearson and Spearman correlation methods.

**Supplemental Figure 1. (A**) Representative changes in *HIF3A* mRNA levels in HUVECs after exposure to hypoxia for the time periods specified were quantified based on expression microarray signals and expressed as fold change relative to normoxic conditions. The data represents mean of 3 biological replicates ± SEM of (\**P*< 0.05, \*\**P*< 0.01 and \*\*\**P*< 0.001 were considered significant). (**B**) *HIF3A silencing in HUVECs exposed to hypoxia increases the expression of HIF1A-AS2. HIF1A*-*AS2* RNA levels were quantified by quantitative real-time PCR and normalized to *RPLP0* rRNA levels and expressed as a fold change over control at the specific time-point of hypoxia. Data represent the mean ± SEM of three replicates (\**P*< 0.05, \*\**P*< 0.01 and \*\*\**P*< 0.001 were considered significant).

## Statements and Declarations

### Author Contributions

Conceived and designed the experiments: R.B., A.B. Performed the experiments: A.B., A.W.W. Analyzed the data: R.B., O.C., P.C. Contributed reagents/materials/analysis tools: R.B. Wrote the paper: J.F.C., L.S., R.B.

## Acknowledgements

We would also like to thank all people that were previously involved in this project, especially Dr. Michał Dąbrowski and Dr. David K. Crossman.

## Data Availability

The accompanying data are included in supplemental materials or available on request.

## Abbreviations

ARE: AU-rich element
BSA: Bovine serum albumin
ECs: Endothelial cells
EPAS1: Hypoxia-inducible factor 2 alpha
GENT2: Gene Expression database of Normal and Tumor tissues 2
GLUT1: Glucose transporter 1
HAEC: Human aortic endothelial cells
HCAEC: Human cardiac artery endothelial cells
HIAEC: Human iliac artery endothelial cells
HIF: Hypoxia-inducible factor
HIF-1: Hypoxia-inducible factor 1
HIF1A: Hypoxia-inducible factor 1 alpha
HIF1A-AS: HIF1A antisense
HIF-2: Hypoxia-inducible factor 2
HIF-3: Hypoxia-inducible factor 3
HMVEC-D: Human dermal microvascular endothelial cells
HMVEC-L: Human lung microvascular endothelial cells
HPAEC: Human pulmonary artery endothelial cells
HPASMC: Pulmonary artery smooth muscle cells
HRE: Hypoxia response element
HUVEC hTERT: Immortalized human umbilical vein endothelial cells
HUVECs: Human umbilical vein endothelial cells
NATs: Natural antisense transcripts
PHD: Prolyl hydroxylase
RPLP0: Ribosomal protein lateral stalk subunit P0
SD: Standard deviation
TSS: Transcription start site
UtMVEC: Human uterine microvascular endothelial cells
UTR: Untranslated region
VEGFA: Vascular endothelial growth factor A
VHL: Von Hippel-Lindau gene

## References

1. Choudhry H, Harris AL (2018) Advances in Hypoxia-Inducible Factor Biology. Cell Metab 27 (2):281–298. doi:10.1016/j.cmet.2017.10.005

2. Wenger RH, Kurtcuoglu V, Scholz CC, Marti HH, Hoogewijs D (2015) Frequently asked questions in hypoxia research. Hypoxia (Auckl) 3:35–43. doi:10.2147/HP.S92198

3. Kaelin WG, Jr., Ratcliffe PJ (2008) Oxygen sensing by metazoans: the central role of the HIF hydroxylase pathway. Molecular cell 30 (4):393–402. doi:10.1016/j.molcel.2008.04.009

4. Kierans SJ, Taylor CT (2021) Regulation of glycolysis by the hypoxia-inducible factor (HIF): implications for cellular physiology. J Physiol 599 (1):23–37. doi:10.1113/JP280572

5. Semenza GL (1999) Regulation of mammalian O2 homeostasis by hypoxia-inducible factor 1. Annual review of cell and developmental biology 15:551–578. doi:10.1146/annurev.cellbio.15.1.551

6. Semenza GL, Roth PH, Fang HM, Wang GL (1994) Transcriptional regulation of genes encoding glycolytic enzymes by hypoxia-inducible factor 1. J Biol Chem 269 (38):23757–23763

7. Jaskiewicz M, Moszynska A, Kroliczewski J, Cabaj A, Bartoszewska S, Charzynska A, Gebert M, Dabrowski M, Collawn JF, Bartoszewski R (2022) The transition from HIF-1 to HIF-2 during prolonged hypoxia results from reactivation of PHDs and HIF1A mRNA instability. Cell Mol Biol Lett 27 (1):109. doi:10.1186/s11658-022-00408-7

8. Hon WC, Wilson MI, Harlos K, Claridge TD, Schofield CJ, Pugh CW, Maxwell PH, Ratcliffe PJ, Stuart DI, Jones EY (2002) Structural basis for the recognition of hydroxyproline in HIF-1 alpha by pVHL. Nature 417 (6892):975–978. doi:10.1038/nature00767

9. Yeh TL, Leissing TM, Abboud MI, Thinnes CC, Atasoylu O, Holt-Martyn JP, Zhang D, Tumber A, Lippl K, Lohans CT, Leung IKH, Morcrette H, Clifton IJ, Claridge TDW, Kawamura A, Flashman E, Lu X, Ratcliffe PJ, Chowdhury R, Pugh CW, Schofield CJ (2017) Molecular and cellular mechanisms of HIF prolyl hydroxylase inhibitors in clinical trials. Chem Sci 8 (11):7651–7668. doi:10.1039/c7sc02103h

10. Hu CJ, Sataur A, Wang L, Chen H, Simon MC (2007) The N-terminal transactivation domain confers target gene specificity of hypoxia-inducible factors HIF-1alpha and HIF-2alpha. Molecular biology of the cell 18 (11):4528–4542. doi:10.1091/mbc.e06-05-0419

11. Ratcliffe PJ (2013) Oxygen sensing and hypoxia signalling pathways in animals: the implications of physiology for cancer. J Physiol 591 (8):2027–2042. doi:10.1113/jphysiol.2013.251470

12. Li C, Jackson RM (2002) Reactive species mechanisms of cellular hypoxia-reoxygenation injury. American journal of physiology Cell physiology 282 (2):C227–241. doi:10.1152/ajpcell.00112.2001

13. Zhang J, Yao M, Xia S, Zeng F, Liu Q (2025) Systematic and comprehensive insights into HIF-1 stabilization under normoxic conditions: implications for cellular adaptation and therapeutic strategies in cancer. Cell Mol Biol Lett 30 (1):2. doi:10.1186/s11658-024-00682-7

14. Slawski J, Jaskiewicz M, Barton A, Koziol S, Collawn JF, Bartoszewski R (2024) Regulation of the HIF switch in human endothelial and cancer cells. Eur J Cell Biol 103 (2):151386. doi:10.1016/j.ejcb.2024.151386

15. Bartoszewski R, Moszynska A, Serocki M, Cabaj A, Polten A, Ochocka R, Dell’Italia L, Bartoszewska S, Kroliczewski J, Dabrowski M, Collawn JF (2019) Primary endothelial cell-specific regulation of hypoxia-inducible factor (HIF)-1 and HIF-2 and their target gene expression profiles during hypoxia. FASEB J 33 (7):7929–7941. doi:10.1096/fj.201802650RR

16. Koh MY, Powis G (2012) Passing the baton: the HIF switch. Trends in biochemical sciences 37 (9):364–372. doi:10.1016/j.tibs.2012.06.004

17. Bartoszewska S, Kochan K, Piotrowski A, Kamysz W, Ochocka RJ, Collawn JF, Bartoszewski R (2015) The hypoxia-inducible miR-429 regulates hypoxia-inducible factor-1alpha expression in human endothelial cells through a negative feedback loop. FASEB J 29 (4):1467–1479. doi:10.1096/fj.14-267054

18. Kochan-Jamrozy K, Kroliczewski J, Moszynska A, Collawn JF, Bartoszewski R (2019) miRNA networks modulate human endothelial cell adaptation to cyclic hypoxia. Cell Signal 54:150–160. doi:10.1016/j.cellsig.2018.11.020

19. Downes N, Niskanen H, Tomas Bosch V, Taipale M, Godiwala M, Vaananen MA, Turunen TA, Aavik E, Laham-Karam N, Yla-Herttuala S, Kaikkonen MU (2023) Hypoxic regulation of hypoxia inducible factor 1 alpha via antisense transcription. J Biol Chem 299 (11):105291. doi:10.1016/j.jbc.2023.105291

20. Serocki M, Bartoszewska S, Janaszak-Jasiecka A, Ochocka RJ, Collawn JF, Bartoszewski R (2018) miRNAs regulate the HIF switch during hypoxia: a novel therapeutic target. Angiogenesis 21 (2):183–202. doi:10.1007/s10456-018-9600-2

21. Chen D, Wu L, Liu L, Gong Q, Zheng J, Peng C, Deng J (2017) Comparison of HIF1A AS1 and HIF1A AS2 in regulating HIF 1alpha and the osteogenic differentiation of PDLCs under hypoxia. Int J Mol Med 40 (5):1529–1536. doi:10.3892/ijmm.2017.3138

22. Reichelt-Wurm S, Knauss L, Strasser B, Scharf M, Holler K, Eggenhofer E, Kretz M, Banas B, Banas MC (2025) Cell-type specific effects of the long non-coding RNA HIF1A-AS3 on HIF1A expression in kidney cells. Scientific reports 15 (1):27876. doi:10.1038/s41598-025-12441-5

23. Zheng F, Chen J, Zhang X, Wang Z, Chen J, Lin X, Huang H, Fu W, Liang J, Wu W, Li B, Yao H, Hu H, Song E (2021) The HIF-1alpha antisense long non-coding RNA drives a positive feedback loop of HIF-1alpha mediated transactivation and glycolysis. Nat Commun 12 (1):1341. doi:10.1038/s41467-021-21535-3

24. Moszynska A, Jaskiewicz M, Serocki M, Cabaj A, Crossman DK, Bartoszewska S, Gebert M, Dabrowski M, Collawn JF, Bartoszewski R (2022) The hypoxia-induced changes in miRNA-mRNA in RNA-induced silencing complexes and HIF-2 induced miRNAs in human endothelial cells. FASEB J 36 (7):e22412. doi:10.1096/fj.202101987R

25. Consortium EP (2012) An integrated encyclopedia of DNA elements in the human genome. Nature 489 (7414):57–74. doi:10.1038/nature11247

26. Kulakovskiy IV, Vorontsov IE, Yevshin IS, Sharipov RN, Fedorova AD, Rumynskiy EI, Medvedeva YA, Magana-Mora A, Bajic VB, Papatsenko DA, Kolpakov FA, Makeev VJ (2018) HOCOMOCO: towards a complete collection of transcription factor binding models for human and mouse via large-scale ChIP-Seq analysis. Nucleic acids research 46 (D1):D252–D259. doi:10.1093/nar/gkx1106

27. Cabaj A, Moszynska A, Charzynska A, Bartoszewski R, Dabrowski M (2022) Functional and HRE motifs count analysis of induction of selected hypoxia-responsive genes by HIF-1 and HIF-2 in human umbilical endothelial cells. Cell Signal 90:110209. doi:10.1016/j.cellsig.2021.110209

28. Rossignol F, Vache C, Clottes E (2002) Natural antisense transcripts of hypoxia-inducible factor 1alpha are detected in different normal and tumour human tissues. Gene 299 (1-2):135–140. doi:10.1016/s0378-1119(02)01049-1

29. Uchida T, Rossignol F, Matthay MA, Mounier R, Couette S, Clottes E, Clerici C (2004) Prolonged hypoxia differentially regulates hypoxia-inducible factor (HIF)-1alpha and HIF-2alpha expression in lung epithelial cells: implication of natural antisense HIF-1alpha. J Biol Chem 279 (15):14871–14878. doi:10.1074/jbc.M400461200

30. Park SJ, Yoon BH, Kim SK, Kim SY (2019) GENT2: an updated gene expression database for normal and tumor tissues. BMC Med Genomics 12 (Suppl 5):101. doi:10.1186/s12920-019-0514-7

31. Jaskiewicz M, Moszynska A, Gebert M, Collawn JF, Bartoszewski R (2023) EPAS1 resistance to miRNA-based regulation contributes to prolonged expression of HIF-2 during hypoxia in human endothelial cells. Gene 868:147376. doi:10.1016/j.gene.2023.147376

32. Ho JJ, Metcalf JL, Yan MS, Turgeon PJ, Wang JJ, Chalsev M, Petruzziello-Pellegrini TN, Tsui AK, He JZ, Dhamko H, Man HS, Robb GB, Teh BT, Ohh M, Marsden PA (2012) Functional importance of Dicer protein in the adaptive cellular response to hypoxia. J Biol Chem 287 (34):29003–29020. doi:10.1074/jbc.M112.373365

33. Balbin OA, Malik R, Dhanasekaran SM, Prensner JR, Cao X, Wu Y-M, Robinson D, Wang R, Chen G, Beer DG (2015) The landscape of antisense gene expression in human cancers. Genome research 25 (7):1068–1079

34. Choi HM, Lee SH, Lee MS, Park D, Choi SS (2021) Investigation of the putative role of antisense transcripts as regulators of sense transcripts by correlation analysis of sense antisense pairs in colorectal cancers. The FASEB Journal 35 (4):e21482

35. Thrash-Bingham CA, Tartof KD (1999) aHIF: a natural antisense transcript overexpressed in human renal cancer and during hypoxia. J Natl Cancer Inst 91 (2):143–151. doi:10.1093/jnci/91.2.143

36. Faridani OR, McInerney GM, Gradin K, Good L (2008) Specific ligation to double-stranded RNA for analysis of cellular RNA::RNA interactions. Nucleic acids research 36 (16):e99. doi:10.1093/nar/gkn445

37. Li L, Wang M, Mei Z, Cao W, Yang Y, Wang Y, Wen A (2017) lncRNAs HIF1A-AS2 facilitates the up-regulation of HIF-1alpha by sponging to miR-153-3p, whereby promoting angiogenesis in HUVECs in hypoxia. Biomed Pharmacother 96:165–172. doi:10.1016/j.biopha.2017.09.113

38. Bartoszewska S, Kamysz W, Jakiela B, Sanak M, Kroliczewski J, Bebok Z, Bartoszewski R, Collawn JF (2017) miR-200b downregulates CFTR during hypoxia in human lung epithelial cells. Cell Mol Biol Lett 22:23. doi:10.1186/s11658-017-0054-0

39. Cayre A, Rossignol F, Clottes E, Penault-Llorca F (2003) aHIF but not HIF-1alpha transcript is a poor prognostic marker in human breast cancer. Breast Cancer Res 5 (6):R223–230. doi:10.1186/bcr652

40. Mineo M, Ricklefs F, Rooj AK, Lyons SM, Ivanov P, Ansari KI, Nakano I, Chiocca EA, Godlewski J, Bronisz A (2016) The Long Non-coding RNA HIF1A-AS2 Facilitates the Maintenance of Mesenchymal Glioblastoma Stem-like Cells in Hypoxic Niches. Cell Rep 15 (11):2500–2509. doi:10.1016/j.celrep.2016.05.018

41. Zolk O, Solbach TF, Eschenhagen T, Weidemann A, Fromm MF (2008) Activation of negative regulators of the hypoxia-inducible factor (HIF) pathway in human end-stage heart failure. Biochem Biophys Res Commun 376 (2):315–320. doi:10.1016/j.bbrc.2008.08.152

42. Vausort M, Wagner DR, Devaux Y (2014) Long noncoding RNAs in patients with acute myocardial infarction. Circ Res 115 (7):668–677. doi:10.1161/CIRCRESAHA.115.303836

43. Jaskiewicz M, Moszynska A, Serocki M, Kroliczewski J, Bartoszewska S, Collawn JF, Bartoszewski R (2022) Hypoxia-inducible factor (HIF)-3a2 serves as an endothelial cell fate executor during chronic hypoxia. EXCLI J 21:454–469. doi:10.17179/excli2021-4622

44. Duan C (2016) Hypoxia-inducible factor 3 biology: complexities and emerging themes. American journal of physiology Cell physiology 310 (4):C260–269. doi:10.1152/ajpcell.00315.2015

45. Ravenna L, Salvatori L, Russo MA (2016) HIF3alpha: the little we know. FEBS J 283 (6):993–1003. doi:10.1111/febs.13572

46. Gebert M, Sobolewska A, Bartoszewska S, Cabaj A, Crossman DK, Kroliczewski J, Madanecki P, Dabrowski M, Collawn JF, Bartoszewski R (2021) Genome-wide mRNA profiling identifies X-box-binding protein 1 (XBP1) as an IRE1 and PUMA repressor. Cell Mol Life Sci 78 (21-22):7061–7080. doi:10.1007/s00018-021-03952-1

47. Bartoszewska S, Kroliczewski J, Crossman DK, Pogorzelska A, Baginski M, Collawn JF, Bartoszewski R (2021) Triazoloacridone C-1305 impairs XBP1 splicing by acting as a potential IRE1alpha endoribonuclease inhibitor. Cell Mol Biol Lett 26 (1):11. doi:10.1186/s11658-021-00255-y

48. Bartoszewski R, Gebert M, Janaszak-Jasiecka A, Cabaj A, Kroliczewski J, Bartoszewska S, Sobolewska A, Crossman DK, Ochocka R, Kamysz W, Kalinowski L, Dabrowski M, Collawn JF (2020) Genome-wide mRNA profiling identifies RCAN1 and GADD45A as regulators of the transitional switch from survival to apoptosis during ER stress. FEBS J 287 (14):2923–2947. doi:10.1111/febs.15195

49. Bartoszewska S, Cabaj A, Dabrowski M, Collawn JF, Bartoszewski R (2019) miR-34c-5p modulates X-box-binding protein 1 (XBP1) expression during the adaptive phase of the unfolded protein response. FASEB J 33 (10):11541–11554. doi:10.1096/fj.201900600RR

50. Livak KJ, Schmittgen TD (2001) Analysis of relative gene expression data using real-time quantitative PCR and the 2(-Delta Delta C(T)) Method. Methods (San Diego, Calif) 25 (4):402–408. doi:10.1006/meth.2001.1262

51. Zerbino DR, Achuthan P, Akanni W, Amode MR, Barrell D, Bhai J, Billis K, Cummins C, Gall A, Giron CG, Gil L, Gordon L, Haggerty L, Haskell E, Hourlier T, Izuogu OG, Janacek SH, Juettemann T, To JK, Laird MR, Lavidas I, Liu Z, Loveland JE, Maurel T, McLaren W, Moore B, Mudge J, Murphy DN, Newman V, Nuhn M, Ogeh D, Ong CK, Parker A, Patricio M, Riat HS, Schuilenburg H, Sheppard D, Sparrow H, Taylor K, Thormann A, Vullo A, Walts B, Zadissa A, Frankish A, Hunt SE, Kostadima M, Langridge N, Martin FJ, Muffato M, Perry E, Ruffier M, Staines DM, Trevanion SJ, Aken BL, Cunningham F, Yates A, Flicek P (2018) Ensembl 2018. Nucleic acids research 46 (D1):D754–D761. doi:10.1093/nar/gkx1098

52. Schodel J, Oikonomopoulos S, Ragoussis J, Pugh CW, Ratcliffe PJ, Mole DR (2011) High-resolution genome-wide mapping of HIF-binding sites by ChIP-seq. Blood 117 (23):e207–217. doi:10.1182/blood-2010-10-314427

53. Tausendschon M, Rehli M, Dehne N, Schmidl C, Doring C, Hansmann ML, Brune B (2015) Genome-wide identification of hypoxia-inducible factor-1 and -2 binding sites in hypoxic human macrophages alternatively activated by IL-10. Biochimica et biophysica acta 1849 (1):10–22. doi:10.1016/j.bbagrm.2014.10.006

54. Lee MC, Huang HJ, Chang TH, Huang HC, Hsieh SY, Chen YS, Chou WY, Chiang CH, Lai CH, Shiau CY (2016) Genome-wide analysis of HIF-2alpha chromatin binding sites under normoxia in human bronchial epithelial cells (BEAS-2B) suggests its diverse functions. Scientific reports 6:29311. doi:10.1038/srep29311

55. Smythies JA, Sun M, Masson N, Salama R, Simpson PD, Murray E, Neumann V, Cockman ME, Choudhry H, Ratcliffe PJ, Mole DR (2019) Inherent DNA-binding specificities of the HIF-1alpha and HIF-2alpha transcription factors in chromatin. EMBO reports 20 (1). doi:10.15252/embr.201846401

56. Krystkowiak I, Lenart J, Debski K, Kuterba P, Petas M, Kaminska B, Dabrowski M (2013) Nencki Genomics Database--Ensembl funcgen enhanced with intersections, user data and genome-wide TFBS motifs. Database : the journal of biological databases and curation 2013:bat069. doi:10.1093/database/bat069

