## Supplementary figures and images for "HIF1A-AS2 Regulates Hypoxia-Driven HIF Isoform Switching in Human Endothelial Cells"

### Figure S1

Supplemental Figure 1

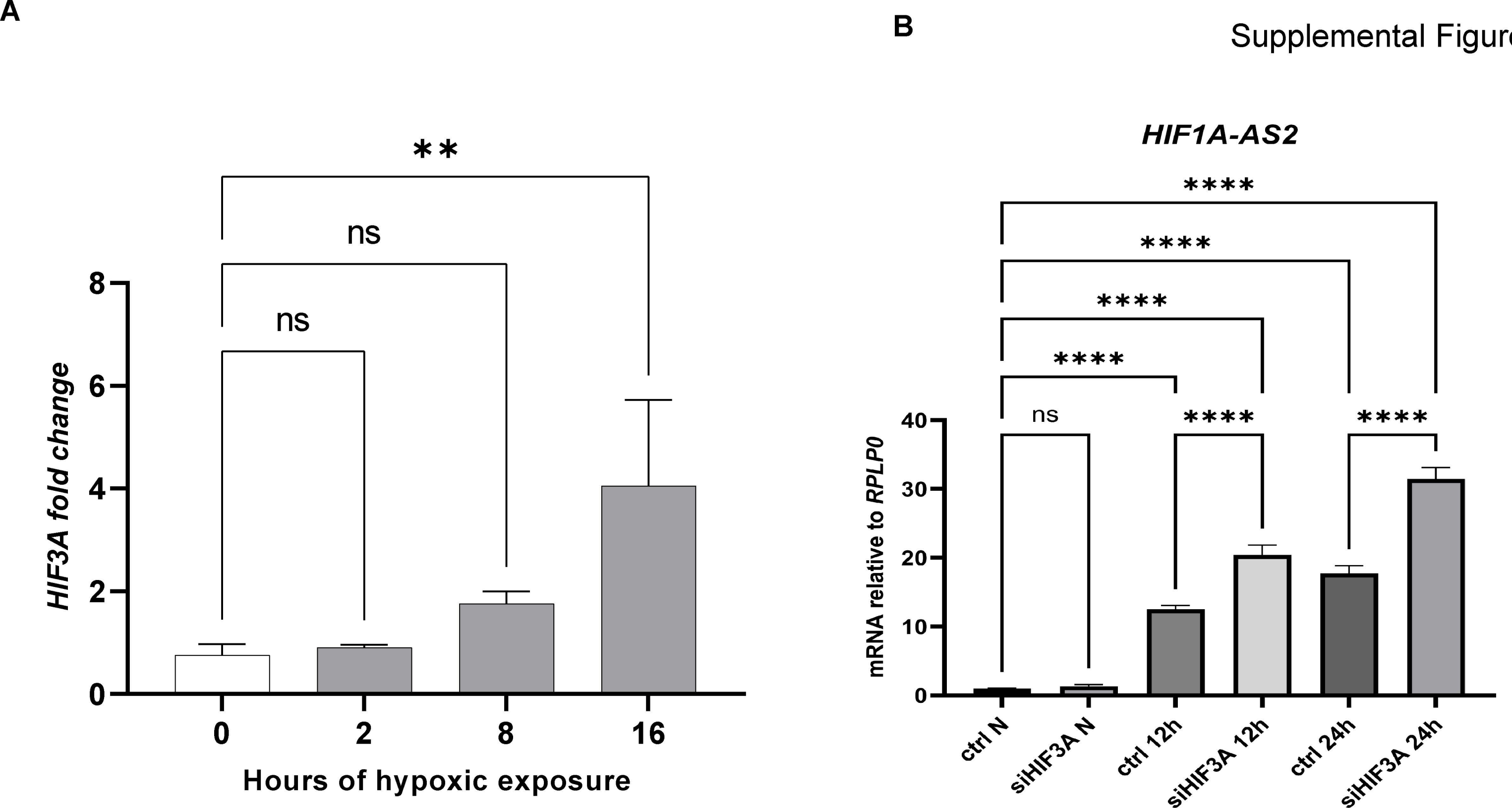

### Figure S2

Uncroped Wb for Figure 1A

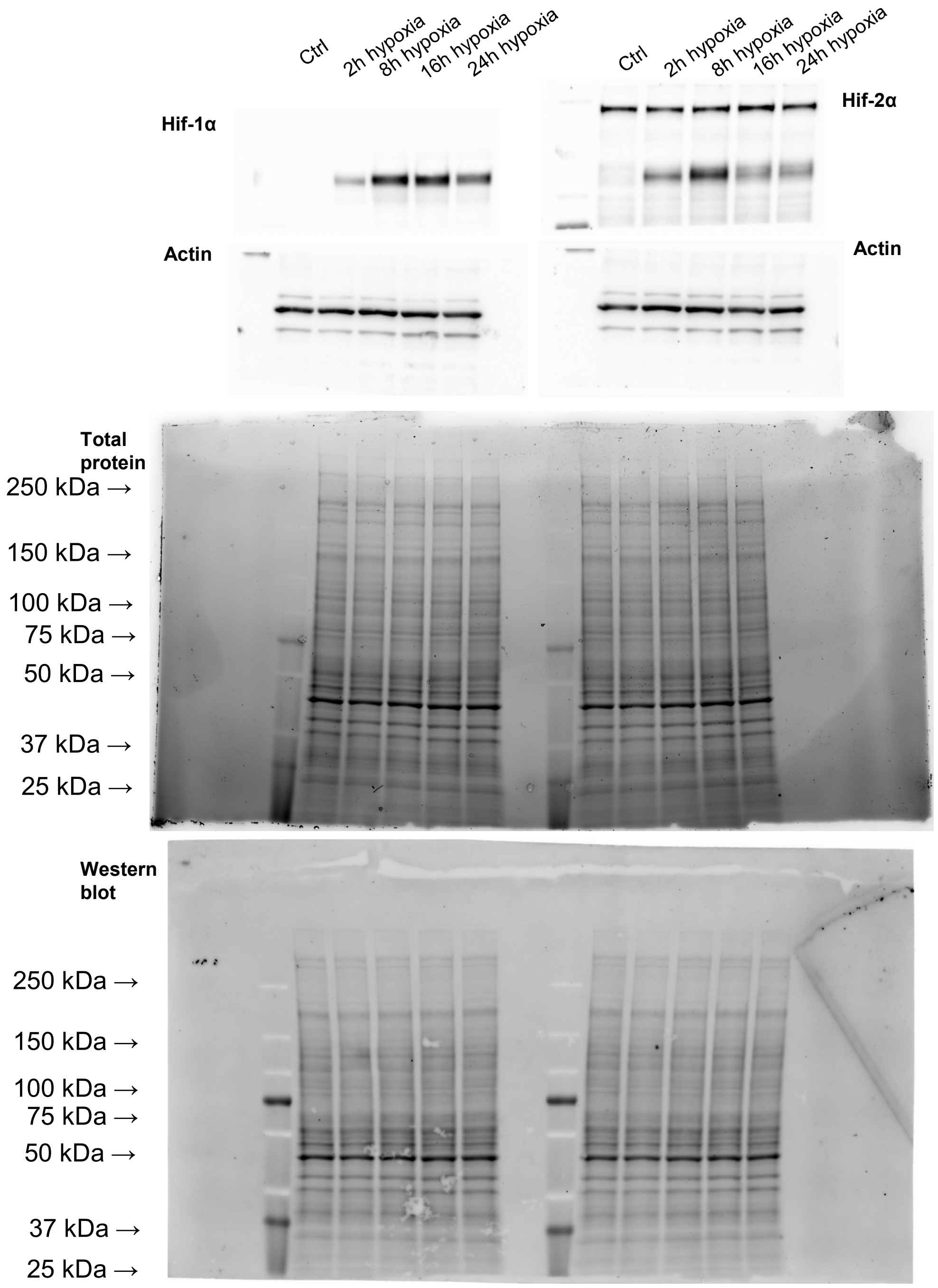

Supplemental Figure 2

#### Uncroped Wb for Figure 6C

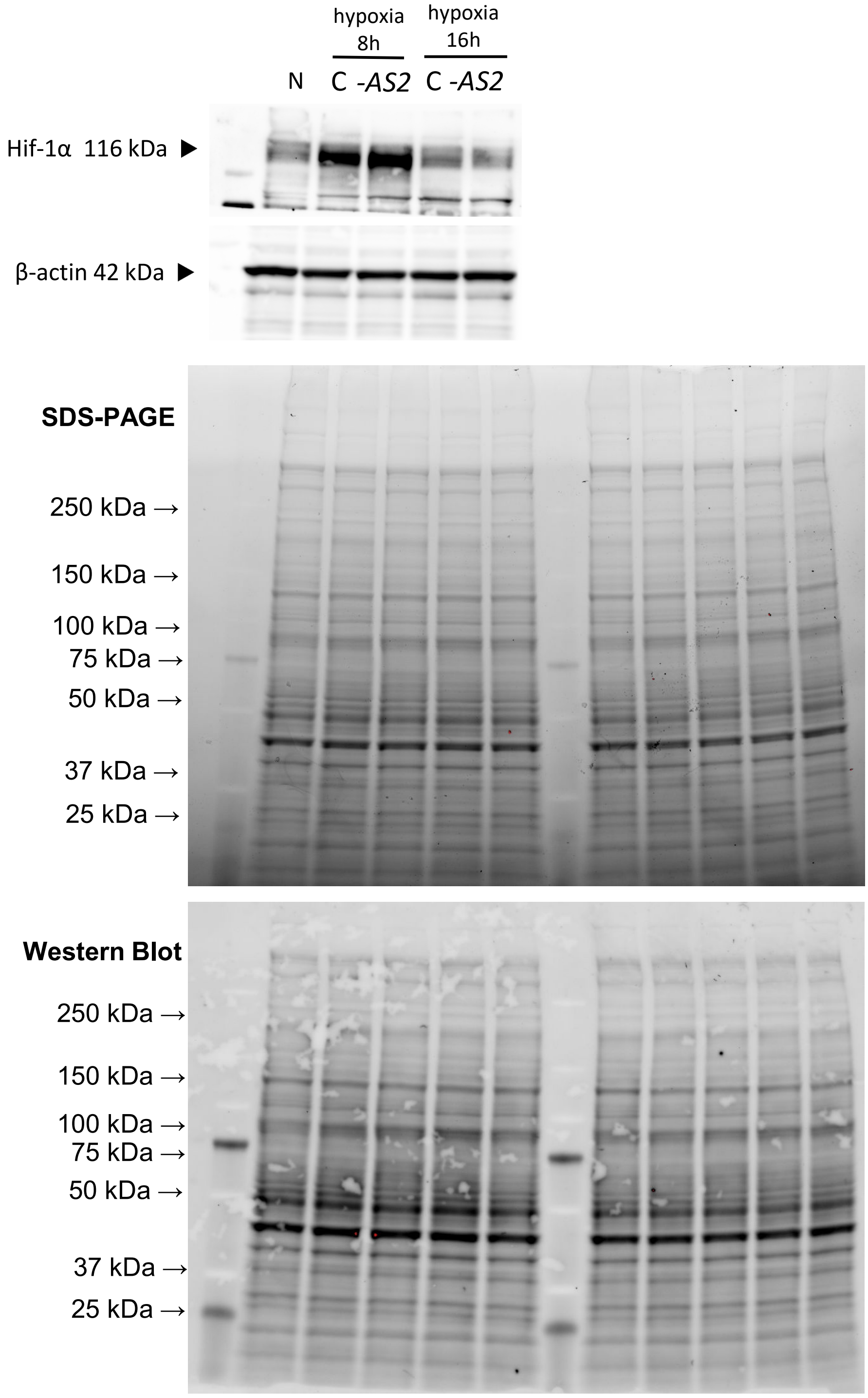
